# Down-regulation of A Human Herpesvirus 1 (HHV-1) MicroRNA in Infected Cells by Goniothalamin Treatment

**DOI:** 10.1101/2021.06.03.447015

**Authors:** Chee Wai Yip, Norefrina Shafinaz Md Nor, Nazlina Ibrahim

**Affiliations:** Department of Biological Science and Biotechnology, Faculty of Science and Technology, Universiti Kebangsaan Malaysia, 43600 Bangi, Selangor, Malaysia

**Keywords:** *Goniothalamus umbrosus*, microRNA, hsv1-miR-H27, Kelch-like 24 protein

## Abstract

Goniothalamin (GTN) has been proven to cause cell cycle arrest and apoptosis in human herpesvirus 1 (HHV-1) infected cells, but interestingly our preliminary transcriptomic analysis revealed other possible modes of action. The data showed that GTN treatment of HHV-1 clinical strain infected cells induced expression of the *KLHL24* gene that encodes the Kelch-like 24 protein (KLHL24), a transcriptional inhibitor of HHV-1 immediate-early and early genes. An miRNA, hsv1-miR-H27, produced by HHV-1 has also been discovered to control the expression of KLHL24. In order to understand the cause of *KLHL24* up-regulation, a time point study was conducted to investigate the effect of GTN on *KLHL24* and hsv1-miR-H27 expression. Through RT-qPCR analysis, we found that HHV-1 down-regulated *KLHL24* significantly (p < 0.05) starting from 12 hpi, while a significant up-regulation (p < 0.05) was observed upon GTN treatment of the infected cells at 4 and 8 hpi. For protein level analysis, significant down-regulation of KLHL24 (p < 0.05) was observed at all time points in HHV-1 infected cells. Intriguingly, treatment with GTN on HHV-1 infected cells showed no significant changes in protein expression compared to cells without any treatment. In addition, the miRNA hsv1-miR-H27 was detected from 16 hpi and treatment with GTN on infected cells showed down-regulation of the miRNA. This was in congruity with the recovery of *KLHL24* down-regulation in GTN treated HHV-1 infected cells, confirming that GTN caused down-regulation of hsv1-miR-H27 that governs the expression of *KLHL24*. This study provides insights that GTN could be a potential multifaceted antiviral.

**Importance:** This study provides evidence that GTN possesses a distinct mode of antivirus against HHV-1 compared to currently available antivirals. Our findings showed that GTN caused the down-regulation of a viral miRNA, which inhibits the expression of a cellular protein known as KLHL24. This protein serves as a transcriptional inhibitor of HHV-1 immediate-early and early genes. The down-regulation of this miRNA lead to the up-regulation of KLHL24 and eventually halted HHV-1 replication. With the previously reported antiviral mechanism and the outcome of this study, GTN is a potential multifaceted anti-HHV-1 agent.

## Introduction

Human herpesvirus 1 (HHV-1) is a highly prevalent member of the Herpesviridae family. The virus has double-stranded DNA and has infected almost 63% of the human population below the age of 50 (James et al. 2020). To date, HHV-1 infections have caused many clinical complications in both immunocompetent and immunocompromised people, including cold sores, acute retinal necrosis, herpes keratitis, esophagitis in transplant patients and encephalitis, which may lead to the death of the infected person (Crimi et al. 2019). Acyclovir (ACV) is an anti-herpetic drug that specifically quells the DNA replication of herpesviruses (Gnann et al. 1983) and was used as a first line treatment to treat primary and recurrent HHV infections. Although this drug is unable to cure latent HHV-1 infections, it has been shown to reduce HHV-1 latency in infected patients (Sawtell et al. 2001). Unfortunately, due to the uncontrolled use of ACV as prophylaxis and in the treatment of immunocompromised patients, resistant strains have emerged (Bacon et al. 2003; Wang et al. 2011). The ability of the virus to confer resistance is caused by mutations, either by addition or deletion, which occur in the *UL23* or *UL30* genes that contribute to the production of different phenotypes of thymidine kinase and DNA polymerase enzymes respectively, thus rendering the drug treatment a failure (Piret & Boivin 2011). Although new drugs such as penciclovir, famciclovir, cidofovir and foscarnet (Superti et al. 2008) have been used to counter the infection of resistant strains, the same restriction has occurred due to the same drug target in the drug design (Piret & Boivin 2011; Wyles et al. 2005). Nonetheless, some of the drugs might pose unwanted side effects for administered patients (Upadhyayula & Michaels 2013). Therefore, this urges the need to search for new anti-HHV agents with novel mechanisms and targets.

Plants are a great source of different groups of secondary metabolites with antiviral properties (Ben-Shabat et al. 2019). The challenges posed by the emergence of antiviral resistant variants have driven scientific communities to search for new antiviral agents from plant sources as some plant extracts exert multifaceted antiviral mechanisms (Álvarez et al. 2011; Schnitzler et al. 2009). *Goniothalamus umbrosus*, also known as ‘kenerak’, is an indigenous plant of Malaysia that possesses various biological properties including antibacterial, antioxidant, anticancer, and antiviral activities (Abdelwahab et al. 2009; Abdul-Wahab et al. 2011). In addition, styrylpyrone derivative (GTN), a bioactive compound isolated from *G. umbrosus* and other species (Jewers et al. 1972; Wiart 2007) has shown potent anti-HHV-1 activity without posing cytotoxic effects on tested cell lines, making it a potential anti-HHV-1 candidate (Md Nor 2011; Moses et al. 2014). Recent findings reported that GTN plays a role in arresting the cell cycle that eventually leads to apoptosis of the infected cells (Md Nor & Ibrahim 2012). Apart from that, the preliminary transcriptomic analysis of Md Nor (2015), using the Next Generation Sequencing (NGS) platform, showed that treatment with GTN on HHV-1 infected Vero cells up-regulated the expression of a cellular protein, namely Kelch-like 24 (*KLHL24*) protein. This protein has been shown to exhibit transcriptional repression of HHV-1 immediate early and early genes (Wu et al. 2013).

Moreover, in order to survive in the host, HHV-1 encodes microRNAs (miRNAs) that control viral gene expression and enable the virus to stay dormant in the host neuron through a process known as latency (Umbach et al. 2008). Latency is maintained by an HHV-1 non-coding transcript which is termed latency associated transcript (LAT). It has been shown that LAT produces miRNAs that can control the gene expression of HHV-1 infected cell protein (*ICP*) *0* and *ICP 4* which are two key transactivators of HHV-1 early gene transcription (Umbach et al. 2008). Besides manipulating viral transcripts, HHV-1 also encodes miRNA that is capable of interfering with host gene expression. The miRNA hsv1-miR-H27 has been proven to cause down-regulation of the host *KLHL24* gene (Wu et al. 2013). However, this miRNA has only been reported in the HHV-1 F strain. No other studies reported the presence of this miRNA in other HHV-1 strains and the presence of miRNA could be viral strain specific (Kim et al. 2012). Therefore, it is necessary to identify the presence of this miRNA in other HHV-1 strains in order to confirm the regulation of *KLHL24* during HHV-1 infection.

This study was conducted to investigate the relationship between *KLHL24* and hsv1-miR-H27 in the absence and presence of GTN by comparing the expression level of both of these elements in HHV-1 infected cells. The outcome of this study will contribute new insights into the anti-HHV-1 properties of GTN and provide strong evidence to develop GTN as an anti-HHV-1 agent with distinct and different modes of action compared to existing chemically synthesised anti-herpetic drugs.

## Materials and Methods

### Cell Culture and Virus Propagation

An African green monkey kidney (Vero) cell line ATCC CCL-81 purchased from American Type Culture Collection (ATCC) was used in this study. The cell line was cultured and maintained in Dulbecco’s modified eagle medium (DMEM) supplemented with fetal bovine serum (FBS) 5% (complete medium) and incubated at 37°C in the presence of 5% CO_2_. A clinical strain of human herpesvirus 1 (HHV-1) was propagated in the Vero cell line. Virus titre determination was performed using plaque assay as described by Blaho et al. (2005) with slight modifications. GTN was extracted according to the method described by Jewers et al. (1972).

### Treatments in Vero Cells

Vero cells were grown until 80% confluency was reached in 25 cm^2^ cell culture flasks (SPL Life Sciences, Korea). Four treatments were given to the Vero cells in each group which included cells mock treated with complete medium, cells treated with 12.5 μM GTN, cells infected with HHV-1 at a multiplicity of infection (MOI) of 1 and HHV-1 clinical strain infected cells treated with 12.5 μM GTN. For treatments involving virus infection, the cells were infected with HHV-1 and 2 hours allowed for adsorption. After the adsorption period, the medium was aspirated to remove unadsorbed viruses and replaced with new complete medium or complete medium containing GTN. Mock treated Vero cells were used as a control. Treatments were given for a specific period (i.e., 4, 8, 12, 16, 18, 20 and 24 hours post infection, hpi) and flasks containing cells with different treatments were flash-frozen in liquid nitrogen prior to RNA isolation to attenuate further gene expression occurring in the cells (Yip et al. 2018).

### RNA Isolation and Reverse Transcription Quantitative Real-Time Polymerase Chain Reaction (RT-qPCR)

Total RNA was isolated using TRIsure (Bioline, USA) and reverse transcribed into cDNA before qPCR was performed. First strand cDNA was synthesised using a Tetro cDNA synthesis kit (Bioline, USA) according to the manufacturer’s protocol. The cDNA of each sample was then diluted 5× prior to qPCR analysis. qPCR was conducted using SensiFast Sybergreen Mastermix (Bioline, USA) using a pair of primers specific to *KLHL24* with annealing temperature of 53°C; forward primer: 5’-TGAGAAGACCACTGTTACACGAGC-3’ and reverse primer: 5’-CCTTGGGGACATCATTTCATTC-3’. For miRNA expression analysis, qPCR was conducted with an annealing temperature of 60°C with a forward primer having a sequence of 5’-CGGGTCTGCATTCAAACACAG-3’ and a reverse primer having a sequence of 5’- CAGACCCCTTTCTCCCCC-3’.

### qPCR Analysis

Absolute quantification was applied to compare the mRNA transcript of *KLHL24* and hsv1-miR-H27 between different treatments. A standard curve was produced by using five dilutions of 10-fold serially diluted *KLHL24* cDNA starting at 200 ng. For hsv1-miR-H27 a standard curve was produced by synthetic oligo synthesised by Integrated DNA Technology (IDT) containing a cDNA sequence of hsv1-miR-H27. Information obtained from the qPCR standard curve was used to estimate the amount of mRNA transcript using the following equation (Ilumina 2010):

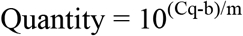

Where,

Cq = Ct values obtained from qPCR result
b = y-intercept of the standard curve
m = gradient of the slope

### Western Blot for Protein Analysis

The organic phase remaining from the previous RNA isolation step for each time point and treatment was used for protein isolation as described by the manufacturer’s protocol. The isolated protein was then subjected to Bradford assay for determination of protein concentration. A total of 50 μg of protein from each treatment was used for sodium dodecyl sulphate polyacrylamide gel electrophoresis (SDS-PAGE) and subsequently Western blotting following the protocol suggested by Mahmood and Yang (2012) with slight modifications. SDS-PAGE was carried out at 75V for 25 minutes followed by 100V for 90 minutes. The separated protein was transferred to a nitrocellulose membrane in Towbin buffer 1× using the Mini Trans-Blot system (Biorad, United States) at 120mA for 120 minutes in ice cold conditions. The primary antibody used for binding to KLHL24 was goat polyclonal antibody IgG anti-KLHL24 at a dilution of 1:500 (Santa-Cruz Biotechnology, Inc., United States). For GAPDH, mouse monoclonal antibody IgG2b anti-GAPDH at a dilution of 1:5000 was used. Secondary antibodies were polyclonal antibody anti-IgG goat (GeneTex, Inc., United States) and anti-IgG1 mouse (Abcam, United Kingdom), both conjugated with horseradish peroxidase (HRP). The substrate for signal detection of the proteins was WesternBright™ Sirius kit (Advansta, Inc., United States).

### Reverse Transcription using a Two-tailed cDNA Synthesis Primer

Two-tailed RT-qPCR was utilised to detect the presence of miRNA hsv1-miR-H27 and also to study the expression profile of this miRNA when different treatments were given to the cells. For detection of hsv1-miR-H27, HHV-1 at MOI 5 was infected to Vero cells for 24 hours before cells were flash-frozen and RNA isolated. For the determination of miRNA expression profile, total RNA in section 2.3 at different time points were used.

First strand cDNA was synthesised using a two-tailed RT primer designed specifically to target hsv1-miR-H27 as described in Androvič et al. (2017). Approximately 500 ng of the total RNA was used for cDNA synthesis using Tetro cDNA synthesis kit with a protocol of 25°C for 45 minutes followed by 85°C for 5 minutes to terminate the reverse transcriptase activity. The sequence for the two-tailed primer used in cDNA synthesis was 5’-GGGTCTGCATTCAAACACAGCTAGAGAACCTAGCTGATCAATTCAAAGAGG-3’.

### Cloning and Sequencing of hsv1-miR-H27 qPCR Product

The qPCR product for hsv1-miR-H27 was first subjected to agarose gel electrophoresis and the band complementary to the size of cDNA was excised and purified using NucleoSpin® Gel and PCR Clean-up (Macherey-Nagel, Germany). The purified product was cloned into T&A™ cloning vector by using T&A™ Cloning Vector (Yeastern Biotech Co., Ltd, Taiwan) and followed by a transformation into *Escherichia coli* E. cloni® 10G strain. Positive transformants were selected using a blue-white screening method. White colonies were picked for colony PCR to further confirm positive transformants. The recombinant plasmid was extracted from the positive transformant using Qiagen® Plasmid Mini (Qiagen, USA) and sequenced (Advanced Innovative Trusted Products & Solutions, AITbiotech Pte. Ltd., Singapore).

## Results and Discussion

### The effect of ***KLHL24*** expression in GTN treated HHV-1 infected or non-infected cells

RT-qPCR analysis showed that the level of *KLHL24* in HHV-1 infected cells was similar to that of the cell only control at time points 4 and 8 hpi but decreased gradually and significantly from 12 hpi (Figure 1). The result is congruent with the study of Wu et al. (2013) where *KLHL24* expression decreased in a time dependent manner coupled with the increase in hsv1-miR-H27. In GTN treated cells, the level of *KLHL24* was higher than the control at all time points. Although treatment of GTN on infected cells increased the expression of *KLHL24* to a level higher than the cell control at earlier time points (4 – 12 hpi), the expression was shown to be slightly lower than the cell control at later time points (16 – 24 hpi). This indicates that the ability of GTN in up-regulating *KLHL24* decreases gradually with time but is still able to maintain similar expression levels as the cell only control.

**Figure 1.**
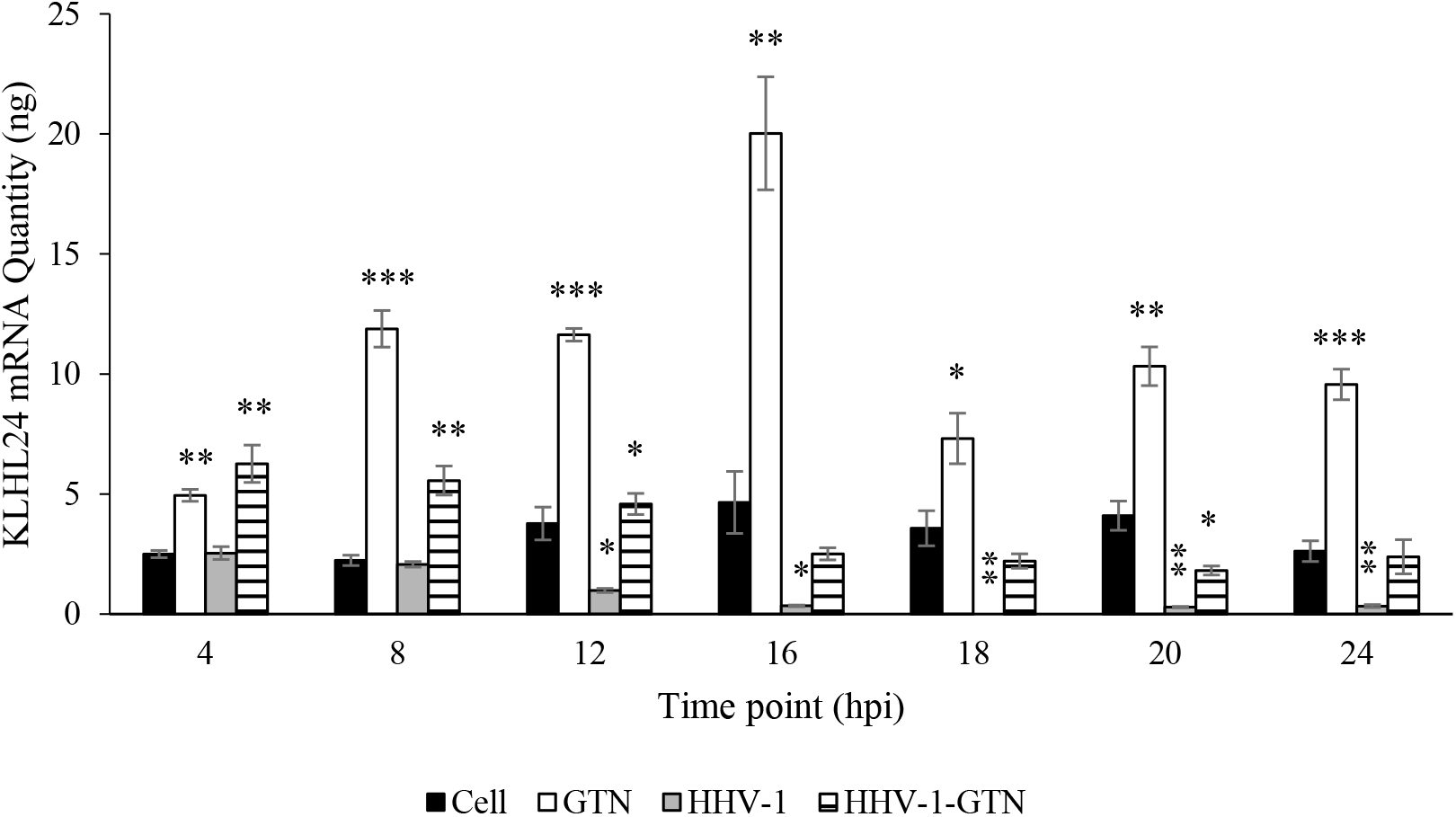
Expression of *KLHL24* analysed by qPCR absolute quantification. The black bar (Cell) represents the amount of *KLHL24* in cell control without any treatment, the white bar (GTN) represents the amount of *KLHL24* in cells treated with 12.5 μM GTN, the grey bar (HHV-1) represents the amount of *KLHL24* in cells infected with HHV-1 at MOI 1, and the horizontal line-patterned bar (HHV-1-GTN) represents the amount of *KLHL24* in HHV-1 infected cells treated with GTN. The data generated was the mean of three replicates together with error bars showing the standard error mean. The significance level p < 0.05 is indicated by *, p < 0.01 is indicated by **, and p < 0.001 is indicated by ***.

Expression in the transcriptional level does not always correlate with the translational level due to post-transcriptional regulation (Edfors et al. 2016). Hence, the protein level expression of KLHL24 was further investigated. Results from Western blot analysis revealed that GTN treatment on non-infected cells showed less than 50% up-regulation of KLHL24 protein level relative to the cell only control and was observed to be not significant at some of the time points (Figure 2). KLHL24 possesses an auto-ubiquitination nature to control its protein level expression if the expression exceeds a normal threshold (Lin et al. 2016). Thus, this property of a negative feedback mechanism might have contributed to the insignificant up-regulation of this protein.

**Figure 2.**
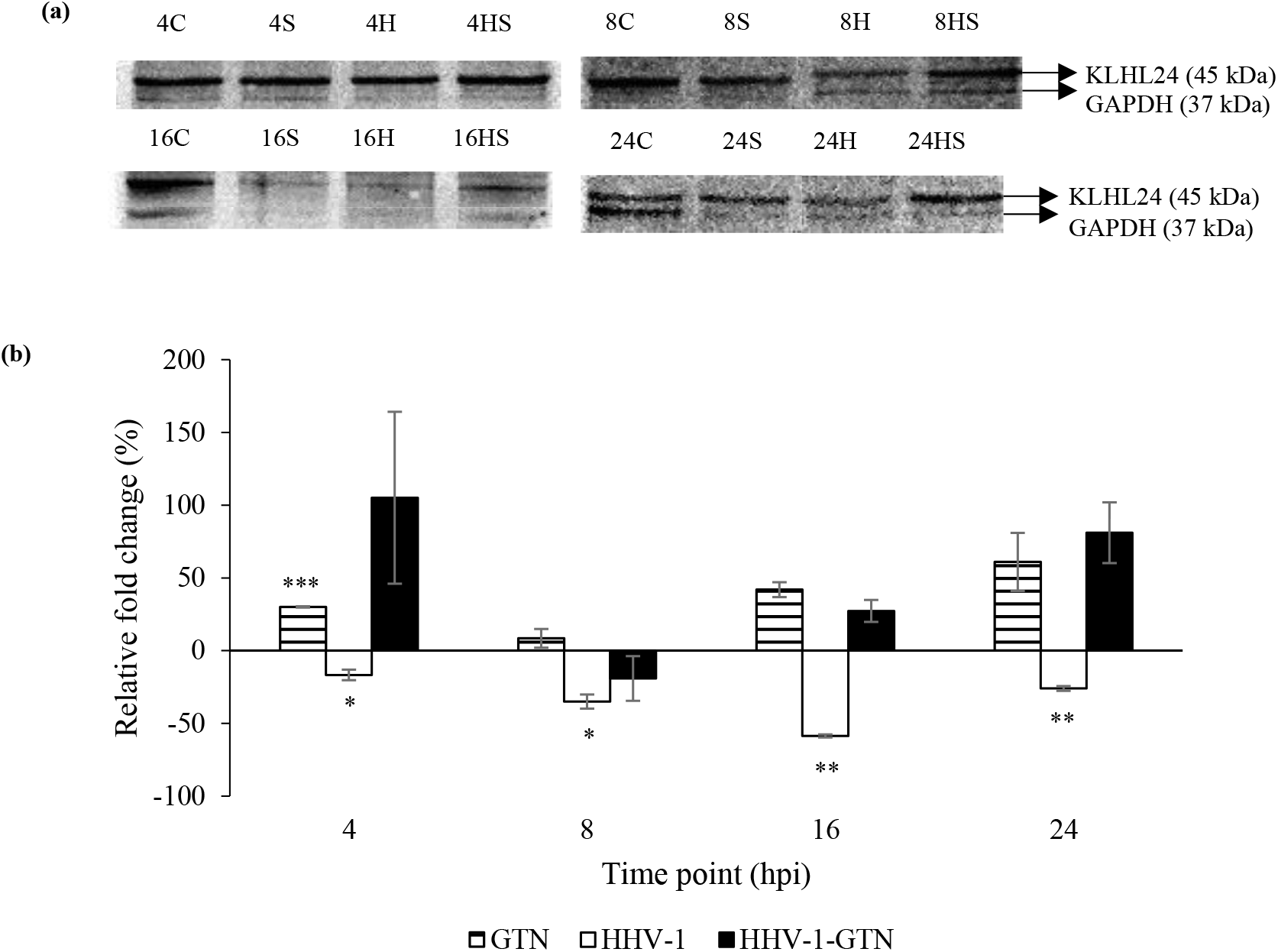
Expression of KLHL24 protein level in different treatments and time points relative to control samples. The horizontal line-patterned bar (GTN) represents Vero cells treated with 12.5 μM GTN, the white bar (HHV-1) represents Vero cells infected with HHV-1 at MOI 1, and the black bar (HHV-1-GTN) represents infected cells treated with GTN. The data generated was the mean of two replicates together with error bars showing the standard error mean. The significance level of p < 0.05 is indicated by *, p < 0.01 is indicated by **, and p < 0.001 is indicated by ***.

In HHV-1 infected cells, the virus significantly down-regulated the expression of KLHL24 at all time points. Interestingly, GTN successfully up-regulated KLHL24 despite the ability of HHV-1, which was shown to inhibit the expression of this protein. As a result, we proved that GTN is able to rescue the down-regulation of KLHL24 caused by HHV-1 infection at both the transcript and protein level.

KLHL24 has been reported as a transcriptional inhibitor for HHV-1 immediate early and early genes, especially *ICP4* (Wu et al. 2013). Additionally, ICP4 functions as a transactivator of HHV-1 early and late genes (Lester & DeLuca 2011). It has been shown that inhibiting the expression of ICP4 will block HHV-1 replication (Wang et al. 2018). Furthermore, treatment with GTN on HHV-1 infected cells led to a down-regulation of ICP4 expression (Md Nor 2015). Therefore, we suggest that the up-regulation of KLHL24 in GTN treated HHV-1 infected cells inhibited ICP4 expression, thereby halting viral replication.

In addition, cell cycle arrest and apoptosis led to an up-regulation of *KLHL24* (Cellai et al. 2009; Hill et al. 2014). GTN has been known to cause cell cycle arrest and apoptosis as the anti-HHV-1 mechanism (Md Nor & Ibrahim 2012). Hence, in this study the up-regulation of KLHL24 is due to the induction of cell cycle arrest and apoptosis by GTN. However, the role of KLHL24 in cell cycle arrest and apoptosis remains elusive to researchers. As KLHL24 is a multifunctional protein (Hedberg-Oldfors et al. 2016; Laezza et al. 2007; Lin et al. 2016), the up-regulation of this protein could lead to the induction of a network of pathways both in the infected host and the virus.

### Detection of hsv1-miR-H27 in HHV-1 Clinical Strain

The miRNA hsv1-miR-H27 was previously reported in HHV-1 F strain (Wu et al. 2013; Du et al. 2015). Nonetheless, the presence of this miRNA is still unknown in other HHV-1 strains and some of the miRNAs are strain specific (Kim et al. 2012). Therefore, the presence of the miRNA needs to be confirmed in the strain used in this study. The result from qPCR produced an amplicon with a size of approximately 70 bp, corresponding to the size of the cDNA which was 68 bp (Figure 3). As it is not practical to sequence a short amplicon, a recombinant plasmid containing the qPCR product was produced for sequencing purposes. The sequence of the recombinant plasmid revealed the presence of 100% sequence similarity of hsv1-miR-H27 in the tested HHV-1 clinical strain. The presence of this miRNA causes down-regulation of *KLHL24* in HHV-1 infected cells. This observation was further confirmed by the miRNA expression study below.

**Figure 3.**
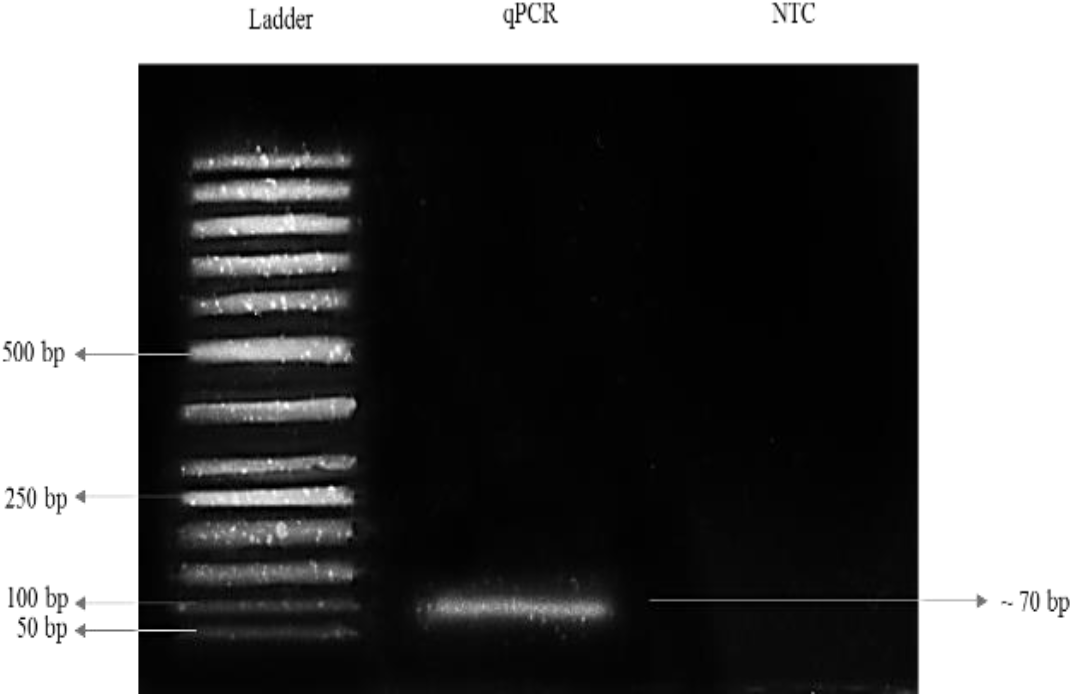
An electropherogram of agarose gel electrophoresis of hsv1-miR-H27 qPCR product. The sample well labelled qPCR represents the qPCR product of hsv1-miR-H27 and the one labelled NTC represents no template control in qPCR. The ladder used in the electrophoresis was GeneRuler^™^ 50 bp DNA ladder (Thermo Scientific^™^, USA).

### The Effect of GTN Treatment on the Expression of hsv1-miR-H27

GTN treatment was shown to cause overexpression of *KLHL24* (Md Nor 2015) and HHV-1 has the ability to control KLHL24 expression by producing hsv1-miR-H27 (Wu et al. 2013). The effect of GTN treatment on the miRNA expression was also tested to identify whether an increase in *KLHL24* expression would cause an increase in the miRNA expression. Our results showed that the miRNA was not detected at an early time point, contradicting the results shown by Wu et al. (2013). This might be due to the different strain of HHV-1 used in the current study compared to the previous study. By 16 hpi, the virus has completed the first round of replication and produced more progeny which produces more miRNA to a level that could be detected at this time point. This was also observed in other studies, which showed the level of miRNA expression was correlated with virus titre (Duan et al. 2012; Flores et al. 2013). Therefore, an increase in miRNA level results in significant down-regulation of *KLHL24* that was observed from 16 hpi (Figure 3 and 4).

**Figure 4.**
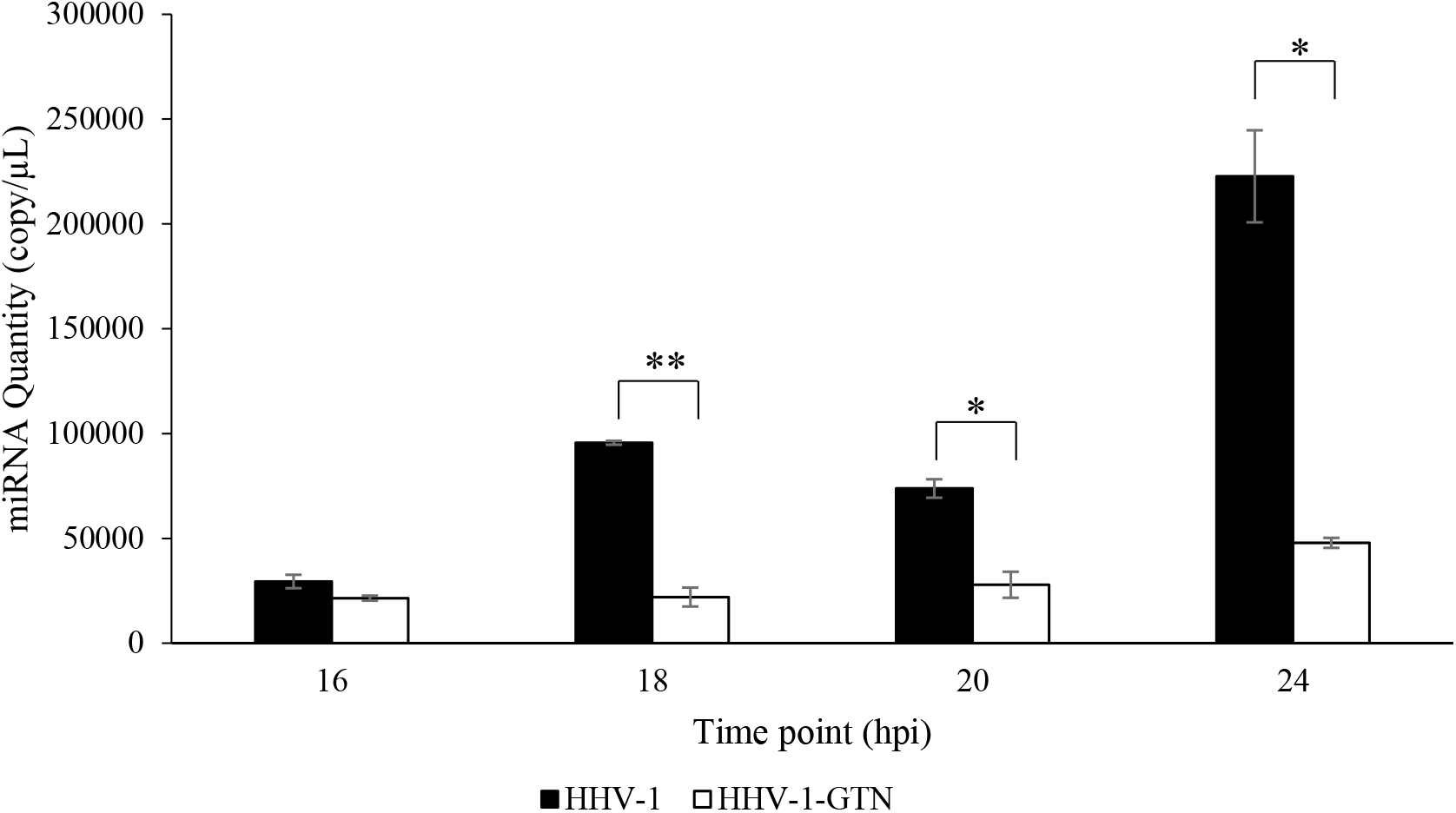
The expression profile of hsv1-miR-H27 in GTN treated and non-treated HHV-1 infected Vero cells. The black bar (HHV-1) represents the copy number of miRNA in HHV-1 infected cells without GTN treatment while the white bar (HHV-1-GTN) represents the copy number of miRNA in HHV-1 infected cells treated with GTN. The data generated was the mean of three replicates together with error bars showing the standard error mean. The significance level of p < 0.05 is indicated by * and p < 0.01 is indicated by **.

Surprisingly, treatment with GTN did not cause up-regulation of the miRNA. Instead, the expression of miRNA was reduced after infected cells were treated with GTN (Figure 4). This result suggested that GTN treatment on the infected cells influenced the expression of this miRNA. Previously, it had been determined that GTN treatment down-regulated the expression of *ICP0*, an immediate early gene of HHV-1 (Md. Nor 2015). The miRNA was predicted to be produced from a precursor, the 3’-untranslated region (UTR) of the *ICP0* gene (Wu et al. 2013). Hence, the down-regulation of hsv1-miR-H27 by GTN treatment was hypothesised to be contributed to by the down-regulation of HHV-1 *ICP0*. Taken together, our results show that GTN caused up-regulation of KLHL24 in HHV-1 infected cells by reducing the expression of hsv1-miR-H27 which is responsible for governing the expression of this gene.

## CONCLUSION

The outcome of this study confirmed that GTN treatment of HHV-1 infected cells caused up-regulation of KLHL24 at gene and protein levels through the down-regulation of hsv1-miR-H27. The down-regulation of hsv1-miR-27 is more likely to be due to the down-regulation of its precursor gene, which affects viral replication.

## ACKNOWLEDGEMENT

The authors would like to acknowledge the Malaysia Ministry of Higher Education for funding this project (FRGS/1/2016/STG04/UKM/02/1). We would also like to thank Prof. Mikael Kubista, Ing. Peter Androvič and the research team for their contribution in designing the two-tailed RT primer required in the RT-qPCR assay of miRNA.

## DECLARATION

The authors declare no conflict of interest in preparing this article.

